# LIM Domain Proteins link molecular and global tension by recognizing strained actin in adhesions

**DOI:** 10.1101/2025.07.16.665189

**Authors:** Stefano Sala, Shreya Chandrasekar, Lee Troughton, Huini Wu, Jordan R. Beach, Patrick W. Oakes

## Abstract

Mechanotransduction is fundamental to cell signaling and depends on force-sensitive adhesion proteins. How these proteins differentiate and integrate their responses to tension remains an open question. We show mechanosensitive LIM domain proteins like zyxin detect global adhesion tension by recognizing strained actin within these structures. In sharp contrast, vinculin localization and intramolecular tension remain unchanged, despite vinculin’s well-documented role in mechanotransduction. This reveals a stark disconnect between molecular tension and global tension in adhesions. We further show tension-dependent localization is specific to LIM domain proteins that recognize strained actin and extends to LIM proteins at cell-cell junctions, suggesting a common mechanotransduction mechanism. Finally, we show zyxin’s tension-dependent adhesion localization stabilizes actin and recruits VASP to promote stress fiber polymerization, identical to its role in stress fiber repair. Our findings reveal a fundamental role for LIM domain protein force-sensing in adhesions and highlight the non-linear connection between molecular and global tension.

## Introduction

Mechanotransduction, like traditional biochemical signaling, modifies cellular behavior across physiology [1–3]. Correspondingly, disruptions in mechanical signaling can contribute to diseases such as cancer, deafness, lung disease, and cardiovascular pathologies [4–6]. The actomyosin cytoskeleton is an essential component of mechanotransduction as it both generates tension and physically connects the internal architecture of the cell to the extracellular environment [7]. Within this cytoskeletal network, multiple molecular-scale mechanisms of force sensing have been discovered [8]. There remains a gap in our knowledge, however, of how tension is propagated across collections of these molecules and how cells sense and respond to macroscopic changes in tension.

Adhesions are critical hubs of mechanical signaling and consist of over a hundred different proteins [9–14]. In focal adhesions (FAs), which couple the cytoskeleton to the extracellular matrix (ECM), these proteins are stratified into distinct layers: a membrane-proximal layer, which includes transmembrane integrins and signaling molecules like FAK, kindlin2 and paxillin; a force-transduction layer, containing adapter proteins like talin and vinculin; and an actin-regulatory layer, where bundles of actin filaments called stress fibers (SFs) terminate and actin-binding proteins such as zyxin, VASP, *α*-actinin and tropomyosins accumulate [15–17]. Many of these individual proteins, including integrins [18], p130Cas [19], FAK [20], talin [21], and vinculin [22] have been shown to change their behavior or function when force is applied directly to the individual protein. Within adhesions, however, these proteins exist as a complex network and display heterogeneous load distributions [23–25]. Unraveling how molecular tension is integrated and contributes to the transmission of FA tension is therefore crucial to understanding FA-mediated mechanotransduction.

The LIM (Lin-11, Isl-1, Mec-3) domain protein zyxin is found in FAs [26], but also plays a key mechanosensing role in coordinating SF repair [27]. When SFs are damaged or begin to fail, tension increases locally along the remaining intact actin filaments, causing strain (i.e. stretch). Zyxin recognizes and binds these strained actin filaments, recruiting VASP and *α*-actinin to promote actin polymerization and crosslinking, thereby stabilizing and repairing the SF [27, 28]. Subsequent studies have identified multiple other LIM domain proteins that similarly recognize strained actin in SFs via their LIM domains, including members of the zyxin, paxillin, testin, Enigma and FHL families [29–32]. Notably, the majority of these mechanosensitive LIM domain proteins also localize specifically to adhesions [33]. How their mechanosensitivity is functionally linked to adhesion signaling, however, remains poorly understood.

Here, we investigated whether zyxin accumulation at FAs is specifically driven by tension and how it contributes to FA signaling. Combining traction force microscopy with photoablation and optogenetic assays, we show that zyxin localization to adhesions directly correlates with FA tension. Surprisingly, both vinculin accumulation and vinculin intramolecular tension within FAs showed no such correlation, despite its established role in mechanotransduction. Instead, this tension-sensitive localization to both FAs, via zyxin and paxillin, and cell-cell adhesions, via TRIP6 and LIMD1, is specific to mechanosensitive LIM domain proteins. We then show that zyxin contributes to tension-sensitive VASP localization to FAs, thereby promoting actin polymerization. Together, these results demonstrate that zyxin recognizes strained actin in adhesions and performs the same function as during SF repair, namely stabilizing filamentous actin to support SF assembly. These data also reveal a fundamental force-sensing mechanism at multiple types of adhesions mediated by the detection of strained actin. Finally, our findings demonstrate that multiple coexisting mechanisms contribute to the conversion of mechanical cues into signaling events at adhesions and provide new insights into how tension propagates across cytoskeletal structures to regulate cellular behavior.

## Results

### Zyxin localization at FAs depends on adhesion tension

Our previous work using machine learning predicted a correlation between zyxin intensity at FAs and traction stresses, suggesting its localization depends on FA tension [34]. To test this hypothesis directly, we used a tightly focused laser to damage a SF upstream of a mature FA, and monitored changes in zyxin intensity and traction stresses at the adhesion (Fig. 1A,B). Photoablation induced a SF strain site and led to local zyxin recruitment (Fig. 1A,C), consistent with previous results [27, 31, 32]. In FAs attached to the damaged SF, we saw a sudden 15-20% drop in zyxin intensity and a simultaneous 30-40% decrease in local traction stresses (Fig. 1D-E; Movie 1), indicating reduced FA tension. This response was specific to adhesions attached to the damaged SF, as adhesions on the opposite side of the cell showed no changes in zyxin intensity or local traction stresses (Fig. 1A-B,D-E).

**Fig. 1.**
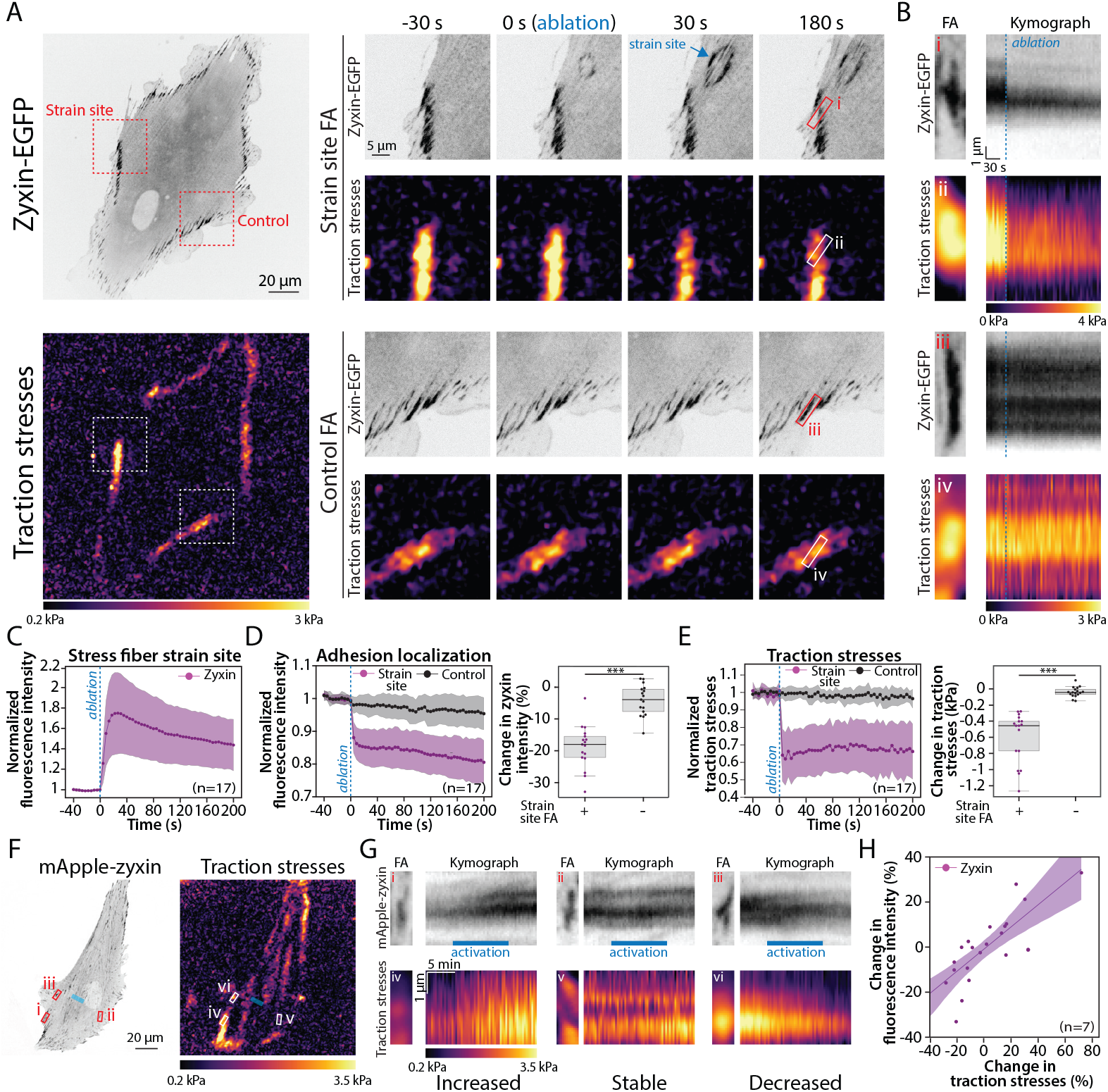
Zyxin localization at FAs scales with adhesion tension. **A)** A representative human foreskin fibroblast (HFF) expressing zyxin-EGFP and the corresponding traction stress map. Insets show zyxin localization and traction stresses over time in the strain site FA and control FA regions (Movie 1). Blue arrow marks zyxin recruitment to SF strain site. **B)** Kymographs of regions marked in (A). **C)** Normalized zyxin intensity *±* SD at SF strain sites. **D-E)** Normalized zyxin intensity (D) and traction stress (E) *±* SD in FAs attached to SFs with photoablation induced strain sites. Boxplots show average changes between the 40s before ablation and the final 40s of each timelapse. **F)** A representative HFF expressing mApple-zyxin and corresponding traction stress map. Blue rectangle marks the optogenetic activation region. **G)** Kymographs of boxed regions in (F), (Movie 2). **H)** Change in zyxin intensity vs. traction stresses post activation. Each point on the scatter plot represents an individual FA. Image intensities in (B) were rescaled only for visualization. *** p *≤* 0.001. n values indicate the number of cells analyzed. Dashed blue lines in (B-E) indicate the ablation timepoint.

Because photoablation only reduces tension, we next investigated how zyxin responds to increased tension at FAs. We used an optogenetic probe to locally activate RhoA, thereby increasing tension in the cytoskeleton [35, 36]. In response to local RhoA activation over a 10 min period, we observed rapid heterogeneous changes in traction stresses across the cell, reflecting the cell’s response to changes in signaling and overall tension (Fig. 1F). We therefore identified regions where traction stresses increased, remained stable, or decreased, and measured changes in zyxin intensity in the corresponding adhesions (Fig. 1G; Movie 2). To isolate tension-dependent responses, we only analyzed adhesions that were morphologically stable. We found that zyxin intensity changes were strongly correlated with traction stress changes, whether assessed at peak response (Fig. 1H), or across individual time points throughout the experiment (Fig. S1A). These results demonstrate that zyxin accumulation at FAs is directly regulated by FA tension.

### Vinculin localization at FAs is independent of adhesion tension

Given the tension-dependence of zyxin localization at FAs (Fig.1), we next examined whether this sensitivity is unique to zyxin. Vinculin plays a key role in mechanotransduction by directly binding actin and other FA proteins such as talin, thereby coupling the actin cytoskeleton to the ECM and promoting force transmission [37] (Fig. 2A). Using the same photoablation approach as above, we simultaneously measured vinculin and zyxin intensity changes in mature adhesions attached to the ablated SF. While zyxin intensity again dropped by ~25% immediately following tension loss, vinculin intensity remained unchanged in the same adhesion (Fig. 2B-D; Movie 3). Control adhesions outside the photoablation region showed no change in either protein’s intensity (Fig. 2B-D). We then used our optogenetic probe to locally activate RhoA and concurrently monitor changes in vinculin intensity and traction stresses (Fig. 2E). In contrast to zyxin, vinculin intensity remained constant despite changes in traction stresses (Fig. 2F-G; Fig. S1B; Movie 2). Together, these results indicate that vinculin localization does not depend specifically on FA tension.

**Fig. 2.**
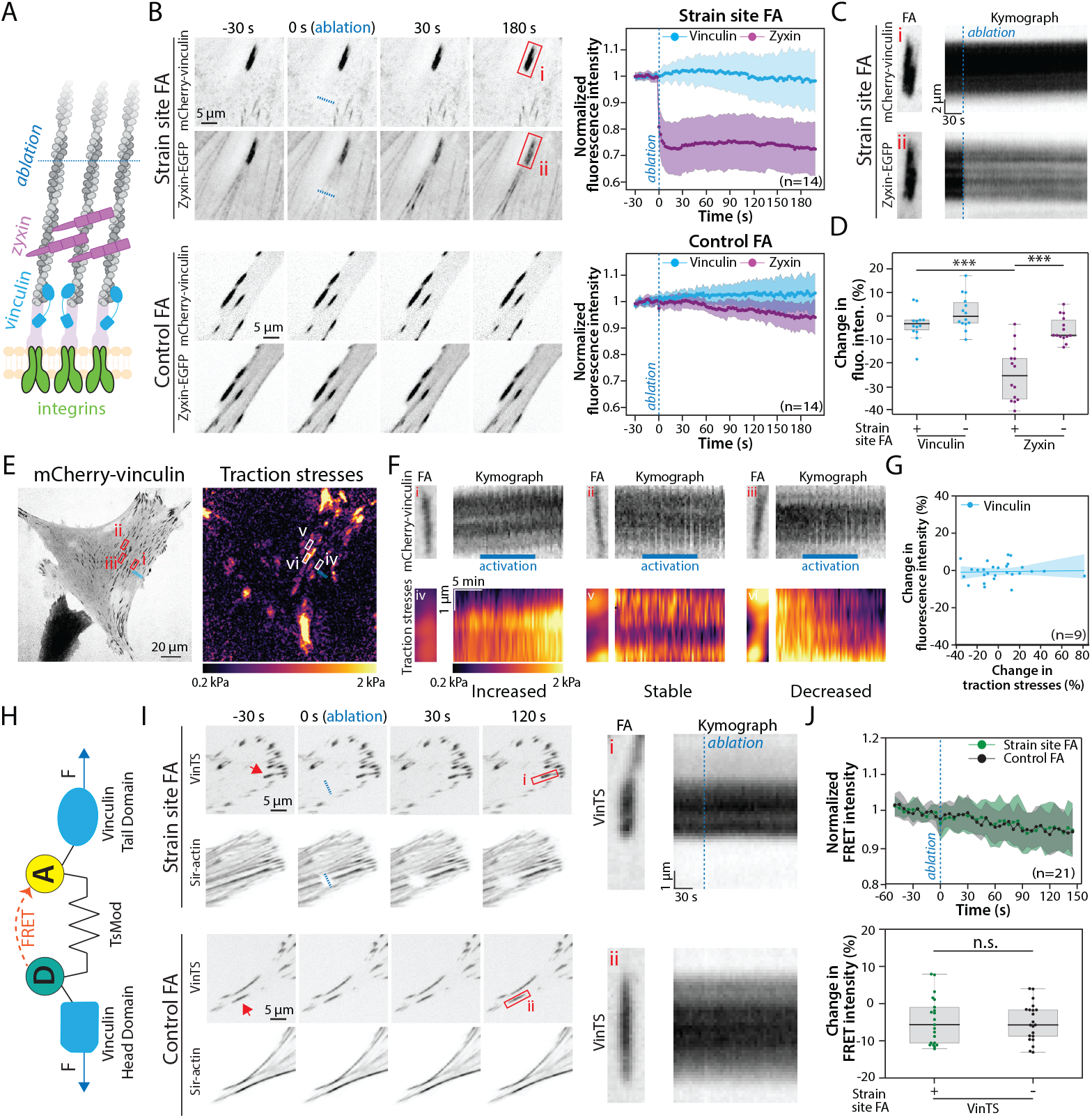
Vinculin localization and intramolecular tension are independent of FA tension. **A)** Schematic showing the relative position of zyxin and vinculin within an FA relative to the ablation region. **B)** Time series of mCherry-vinculin and zyxin-EGFP in FAs of HFFs attached to SFs with strain sites or control (Movie 3). Normalized fluorescence intensity *±* SD of vinculin and zyxin in FAs. **C)** Kymographs of the red boxes in (B). **D)** Boxplots showing change in average zyxin or vinculin intensities between the 30s before ablation and the final 30s of each timelapse. **E)** A representative HFF expressing mCherry-vinculin and corresponding traction stress map. Blue rectangle marks the optogenetic activation region. **F)** Kymographs of boxed regions in (E), (Movie 2). **G)** Change in vinculin intensity vs. traction stresses post activation. Each point on the scatter plot represents an individual FA. **H)** Schematic of the FRET-based vinculin tension sensor (VinTS). **I)** Mouse embryonic fibroblasts (MEFs) stably expressing VinTS showing its localization over time in the strain site and control FAs (Movie 4). Red arrows indicate the adhesions used for analysis. Red boxed regions indicate the kymographs on the right. **J)** Normalized FRET intensity *±* SD in strain site and control FAs. Boxplots show the change in FRET intensity between the 25s before ablation and last 25s of each timelapse. *** p *≤* 0.001. n values indicate the number of cells analyzed. Dashed blue lines in B, C, I, and J indicate the ablation timepoint.

### Vinculin intramolecular tension is independent of FA tension

Previous studies have established that vinculin undergoes conformational changes when subjected to mechanical load [22, 38]. We therefore explicitly tested whether reducing FA tension influences intramolecular tension across vinculin using a well-characterized FRET-based vinculin tension sensor (VinTS) [22, 25]. As tension on vinculin increases, the elastic linker stretches, increasing the distance between the fluorophores and thereby decreasing FRET intensity (Fig. 2H) [22, 25]. To minimize interference from endogenous vinculin, we knocked down its expression (~80% reduction; Fig. S1C), and stably re-expressed the VinTS construct (Fig. S1D). Surprisingly, FRET intensity in mature stable adhesions remained unchanged following photoablation of the attached SF, similar to control adhesions (Fig. 2I-J; Movie 4). To confirm that our setup could detect changes in FRET intensity, we treated cells with 50 *µ*M Y-27632, a ROCK inhibitor known to reduce cytoskeletal tension and increase FRET in VinTS [22, 39]. Consistent with previous results, we saw a ~10% increase in FRET intensity in FAs following Y-27632 treatment, but not in the media control (Fig. S1E-F). We thus conclude that intramolecular tension on vinculin does not directly correlate with FA tension, and that multiple mechanisms of tension sensing coexist within FAs.

### Tension-dependent FA localization is specific to mechanosensitive LIM domain proteins

Building on our finding that zyxin and vinculin respond differently to changes in adhesion tension (Fig. 2), we asked whether this behavior is specific to LIM domain proteins. To test this, we first examined paxillin, a LIM domain-containing FA protein that, like zyxin, binds strained actin [29]. When we photoablated a SF attached to an adhesion expressing both zyxin and paxillin, their fluoresence intensities decreased by ~20% and ~15%, respectively, while control adhesions remained unaffected (Fig. 3A; Movie 5). In contrast, the FA protein kindlin2, which binds paxillin and clusters integrins during adhesion formation but lacks LIM domains [40], showed no significant change in localization (Fig. 3B; Movie 5). To test whether this response was specific to strain-sensitive LIM domain proteins, we tested the non-mechanosensitive LIM domain protein PINCH1 (also known as LIMS1) [30, 31] and again observed no significant localization change (Fig. 3C; Movie 5). Together, these results indicate that the tension-dependent localization of LIM domain proteins to FAs is specific to those capable of recognizing strained actin filaments.

**Fig. 3.**
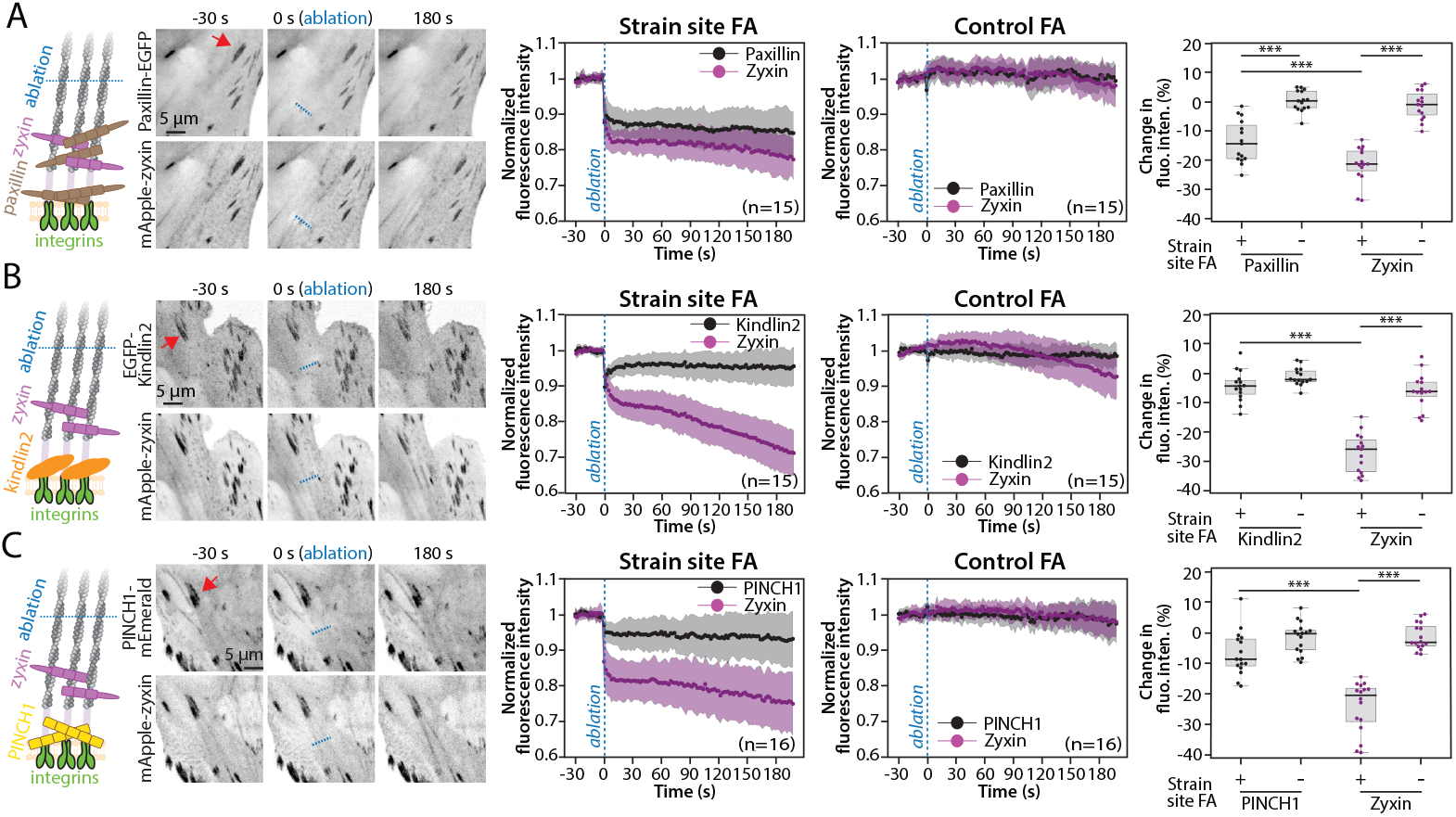
Strain-sensitive LIM domain proteins localize to focal adhesions in a tension-dependent manner. **A-C)** Photoablation experiments in HFFs co-expressing mApple-zyxin and either paxillin-EGFP (A), EGFP-Kindlin2 (B), or PINCH1-mEmerald (C) (Movie 5). Schematics illustrate the relative positions of the proteins in relation to zyxin and the ablation site. Plots display the average FA intensity *±* SD of the proteins of interest. Boxplots indicate the relative change between the average pre-ablation intensity and the final 30s of the movie. Red arrows indicate the adhesions used for analysis. *** p *≤* 0.001. n values indicate the number of cells analyzed. Dashed blue lines indicate the ablation timepoint.

### Trip6 and LIMD1 localization at cell-cell junctions is tension dependent

Analogous to FAs, cell-cell junctions are protein complexes that transmit cytoskeletal tension across the cell membrane to neighboring cells. Previous studies using pharmacological myosin inhibitors have suggested that certain LIM domain proteins, including TRIP6 and LIMD1, are recruited to cell-cell junctions in a tension-dependent manner [41, 42]. To directly test this hypothesis, we adapted our photoablation assay to investigate whether the localization of E-cadherin, TRIP6 and LIMD1 to cell-cell junctions depends on tension within the actin cytoskeleton, using MCF10A epithelial cells. To minimize junction movement, photoablations were performed simultaneously on both sides of the junction. Following photoablation, E-cadherin intensity showed a small (~5%) reduction, likely due to a combination of photobleaching and changes in z-position during imaging (Fig. 4A; Movie 6). In contrast, both Trip6 and LIMD1 intensities showed significantly larger decreases of ~25% and ~15%, respectively, at junctions connected to the ablated SFs (Fig.4B-C; Movie 6). These results indicate that, similar to zyxin and paxillin in FAs, Trip6 and LIMD1 localization to cell-cell adhesions is tension dependent.

**Fig. 4.**
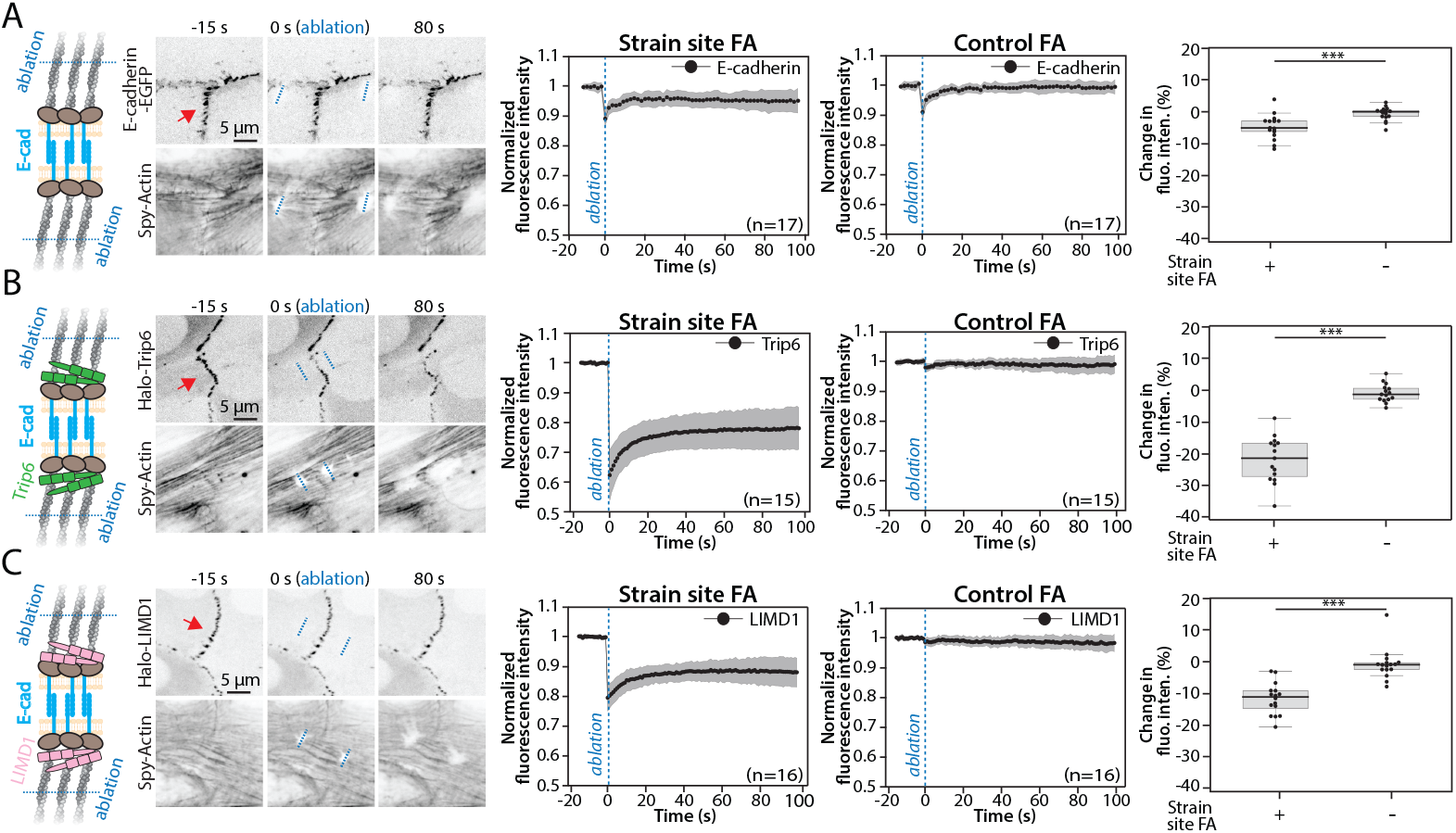
Strain-sensitive LIM domain proteins localize to cell-cell adhesions in a tension-dependent manner. **A-C)** Photoablation experiments in MCF10As expressing E-cadherin-EGFP (A), Halo-Trip6 (B) or Halo-LIMD1 (C) (Movie 6)). Schematics illustrate the relative positions of the proteins in relation to E-cadherin and the ablation site. Plots display the average adhesion intensity *±* SD of the proteins of interest. Boxplots indicate the relative change between the average pre-ablation intensity and the final 15s of the movie. Red arrows indicate the adhesions used for analysis. *** p *≤* 0.001. n values indicate the number of cells analyzed. Dashed blue lines indicate the ablation timepoint.

### Tension-dependent localization of zyxin enables VASP-mediated actin polymerization at FAs

While our results indicate that zyxin recognizes strained actin in FAs, the functional implications of this behavior remain unclear. Recent in vitro work recapitulating zyxin SF repair demonstrated that zyxin recruits VASP to SF strain sites, promoting the polymerization of new actin filaments [28]. If zyxin performs a similar role in FAs, this raises two hypotheses: (1) VASP localization to adhesions is tension dependent, and (2) the loss of zyxin in FAs impairs actin polymerization at these sites. To test the first hypothesis, we expressed VASP in WT and zyxin KO MEFs and photoablated a SF while monitoring VASP intensity in the attached adhesion. In WT cells, VASP intensity steadily declined by ~25% following photoablation (Fig. 5A; Movie 7). In contrast, VASP intensity remained unchanged in zyxin KO cells in response to the same perturbation. These results indicate that zyxin recruits VASP to adhesions in a tension-dependent manner, similar to its role in SF strain sites.

**Fig. 5.**
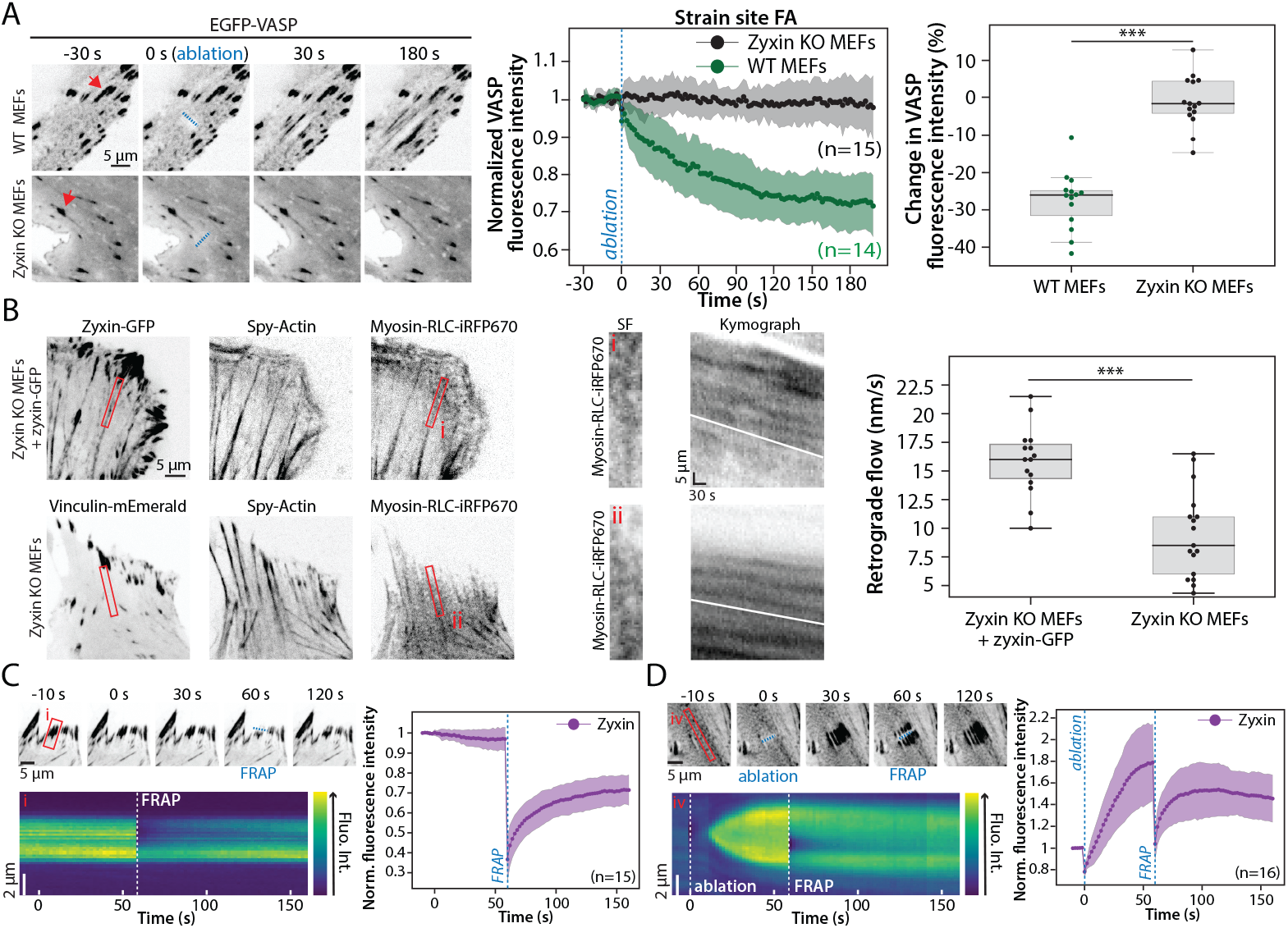
Tension-dependent localization of zyxin enables VASP-mediated actin polymerization at FAs. **A)** WT and zyxin KO MEFs expressing EGFP-VASP (Movie 7). Red arrows mark the adhesions used for analysis. Plots show normalized VASP intensity *±* SD. Boxplots show change in average VASP intensity between the 30s before ablation and final 30s of each movie. **B)** zyxin KO MEFs expressing GFP-zyxin or vinculin-mEmerald, in addition to myosin-RLC-iRFP670 for quantification of actomyosin retrograde flow. Kymographs show myosin RLC signal from red-boxed regions. Boxplots show average retrograde flow velocities per cell in WT and KO MEFs. **C-D)** Representative images and kymographs (red boxes) of MEFs expressing zyxin-GFP with FRAP and ablation regions indicated by the blue dashed lines. Plots display normalized zyxin intensity over time in the targeted regions. *** p *≤* 0.001. n values indicate the number of cells analyzed.

To test the second hypothesis, we measured actomyosin retrograde flow originating from FAs in zyxin KO MEFs +/zyxinGFP. Kymograph analysis revealed an average retrograde flow rate of 15-20 nm/s in MEFs with zyxin, consistent with previous findings [43], compared to ~10 nm/s in zyxin KO MEFs (Fig.5B). These data indicate that zyxin promotes actin polymerization at FAs, suggesting it performs the same functional role as in SF strain sites.

### Zyxin exchange dynamics in SF strain sites and FAs are similar

If zyxin has a similar role in FAs and SF strain sites, its dynamics in strain sites should be similar to its dynamics in FAs. We therefore performed FRAP experiments and confirmed that zyxin turns over rapidly in FAs, with a *t*_1_*/*2 on the order of 10s (Fig. 5C; [44]). Next, we performed similar experiments at SF strain sites. First, we measured zyxin accumulation at these strain sites to determine the appropriate FRAP timepoint and found that zyxin intensity peaked approximately 60s post ablation (Fig. S2A). We then validated our FRAP settings by bleaching an unstrained SF segment, confirming that the light intensity used did not induce strain sites and thus spur additional zyxin recruitment (Fig. S2B). Finally, we combined both approaches: we first induced a strain site in the SF and then performed FRAP on a central region in that site (Fig. 5D). Zyxin showed rapid recovery at strain sites, indicative of fast turnover, with rates comparable to those we observed in FAs. Together, these results demonstrate that zyxin rapidly exchanges in both FAs and SF strain sites, recognizing strained actin and promoting actin polymerization through VASP recruitment (Fig. 6).

**Fig. 6.**
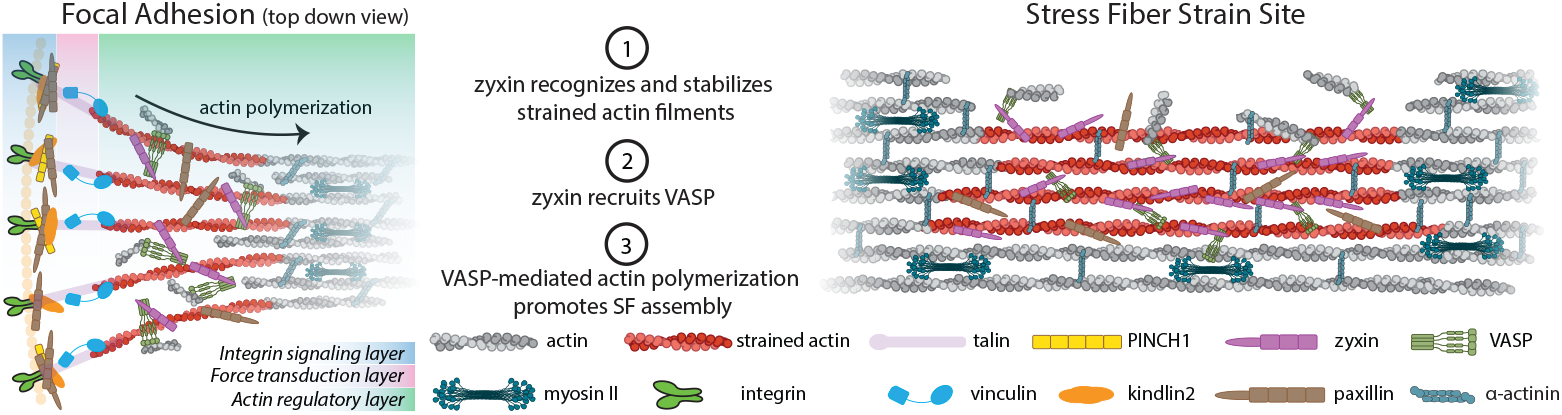
Zyxin performs the same functions at FAs and SF strain sites. At both FAs and strain sites, zyxin recognizes strained actin and stabilizes it while it recruits VASP to promote actin polymerization.

## Discussion

Our findings demonstrate that mechanosensitive LIM domain proteins specifically recognize strained actin at adhesions, providing a molecular basis for their tension-dependent localization to these structures. Strained actin is typically found in SFs at sites of damage or failure, where the intact filaments locally experience higher load and likely stretch slightly [27, 45]. Zyxin binds these strained filaments through its C-terminal LIM domains and recruits VASP and *α*-actinin through N-terminal motifs to facilitate rebuilding the SF [27, 28]. Our data indicate that a mechanically similar process occurs naturally in FAs (Fig. 5). During SF assembly at FAs, newly polymerized actin filaments likely also experience a higher load [46], thereby causing strain until they are fully crosslinked into a SF bundle. In this context, we propose that the role of zyxin is not limited to mediating SF repair, but reflects a broader function in actin strain-sensing and stabilization during de novo SF assembly in FAs (Fig. 6).

The striking difference in response to changes in tension we see between zyxin and vinculin (Fig. 2) demonstrates the presence of multiple mechanosensing mechanisms acting in parallel. These results are consistent with previous work that identified increased exchange kinetics for zyxin in FAs with decreased tension, but not for vinculin [44]. More broadly, previous studies have shown that within an adhesion, proteins exhibit highly heterogeneous properties. For example, single molecule tracking of integrins reveals that only a subpopulation of integrins in an FA is bound to ECM at a given time [47]. The load on these integrins is also heterogeneously distributed across the FA [23, 48]. Similarly, tension sensors inserted into talin reveal that only ~40-60% of talin molecules are loaded at any given time [24], while vinculin itself displays a non-uniform distribution of load within the FA [22, 25]. This high degree of heterogeneity of protein behaviors in FAs indicates that there are limitations connecting single-molecule measurements to adhesion network measurements, such as the traction stresses we measure here. In this case, our data shows that zyxin accumulation serves as a more reliable proxy of FA tension than vinculin.

This is not to suggest that vinculin or other FA proteins are not mechanosensitive. Because the tension changes we induce are the result of perturbations on the SF upstream of the FA, the local SF environment in adhesions where vinculin is located likely remains constant. We thus speculate that the tension-dependent conformational change of actin is sufficient to alter binding of zyxin, but is not large enough to displace or alter tension on vinculin. In contrast, pharmacological treatments with Y-27632 or blebbistatin reduce global tension, but they also severely disrupt actin cytoskeleton architecture [49]. This could contribute to the large number of proteins that are identified as mechanoresponsive to pharmacologic perturbations of tension in proteomic screens [10, 11, 50]. Our results therefore illustrate the importance of architecture in conjunction with tension in regulating mechanosensitive interactions, and highlight the non-linear connection between molecular tension and FA tension. By inducing acute tension changes and focusing on mature, stable adhesions that are not assembling or disassembling, our approach minimizes the effects from architectural changes, allowing us to isolate these different tension-dependent mechanisms.

Interestingly, we see a reduced tension-sensitive response of paxillin compared to zyxin (Fig. 3A). This could be explained by two separate, non-exclusive, hypotheses. First, it is possible that paxillin and zyxin sense different magnitudes of strain. While our data does not explicitly address this hypothesis, it is appealing as it would enable the cell to trigger different mechanotransduction pathways depending on the magnitude of local strain in the adhesion. Alternatively, a reduction in paxillin could indicate that there are different subpopulations of paxillin in the adhesion. In addition to actin, paxillin binds multiple proteins, including integrins [51], FAK [52], and kindlin [40] which are all located proximal to the membrane. While a reduction in tension could modulate paxillin binding to actin, these other interactions are likely tension-independent. Neither of these hypotheses are restricted to paxillin and we suspect that a thorough cataloging of additional LIM proteins would reveal a range of responses. Resolving these hypotheses is an area of future interest, but we can safely conclude that force sensing occurs across the entire adhesion architecture.

While zyxin clearly facilitates SF assembly, whether in SF strain sites or at adhesions, the role of other mechanosensitive LIM domain proteins remains unclear. Their multimodular structure allows them to scaffold many interaction partners, potentially triggering multiple mechanotransduction pathways [33, 53]. Perhaps their most straightforward role is stabilizing strained actin filaments throughout the cell as these filaments are exposed to an array of different forces [54, 55]. This behavior could sequester LIM domain proteins from the nucleus, thereby regulating their transcriptional activity [56, 57], or even contribute to cell-cycle control [58]. In this scenario, changes in cell morphology that reduce the number of adhesions would also free additional LIM domain proteins to relocalize and alter downstream signaling [59, 60]. Subcellularly, force sensing at individual adhesions could also aid in establishing asymmetries in tension needed to guide migration [61], consistent with recent work showing that zyxin is required for durotaxis [62]. Thus by both direct and indirect mechanisms, and likely in parallel, LIM domain proteins at adhesions contribute to mechanotransduction.

Importantly, while our results focus largely on the behavior of zyxin at FAs, our Trip6 and LIMD1 results indicate that the same interactions are likely at play in cell-cell adhesions (Fig. 4). Mechanistically, the situations are similar - actin is polymerized under tension and crosslinked into a network by a collection of adhesion proteins [63]. While TRIP6 and LIMD1 are not known to contribute to actin polymerization or repair, they sequester LATS1/2 kinases at adhesions, keeping them from phosphorylating, and thereby activating, YAP as part of the Hippo signaling pathway [41, 42]. Further investigation will be needed to explore the mechanistic roles of these other LIM domain proteins in these structures.

Finally, our data support a common mechanism of tension sensing and position LIM domain proteins as a broad class of mechanosensors that recognize strained actin filaments. How strained and unstrained actin layers, along with the many individual mechanosensitive proteins at adhesions, integrate to coordinate signal transduction remains an open question. To fully parse this behavior we need a better census of the number and organization of components and deeper understanding of their parallel responses to local changes in tension. There is also no reason to believe that this strain sensing behavior is limited to adhesions or SF strain sites. Strained actin networks, and probably several LIM domain proteins, are also likely to be found in the cortex, the nucleus, and the contractile ring, for example. This points to broader applications of force-sensing operating across cytoskeletal compartments, each potentially engaging a distinct subset of LIM domain proteins that convert mechanical signals into cytoskeletal remodeling and signaling, thereby maintaining tensional homeostasis in cells.

## Limitations of the study

To date, zyxin remains the only LIM domain protein whose ability to bind strained actin has been reconstituted in purified systems [28, 30, 31]. While our results are indicative of other LIM domain proteins also preferentially binding strained actin, we cannot definitively rule out additional protein-protein interactions contributing to their relocalization [53]. Our work primarily focuses on FAs, but our data in cell-cell adhesions suggest that similar mechanical mechanisms occur at these sites. The specific contributions of Trip6 and LIMD1, or even other LIM domain proteins, to actin polymerization at these sites, however, remains to be clarified. Finally, because of challenges with live-imaging of the FRET constructs, our FRET measurements report only relative changes and not absolute values of tension or FRET efficiency. The lack of change in intensity in these measurements, however, indicates that the conclusion would be the same.

## Materials and Methods

### Expression vectors and molecular cloning

cDNA encoding zyxin was cloned into the mApple-C1 vector (Addgene plasmid #54631) as previously described ([34]). To generate the LV-EGFP-SspB-LARG vector, the LV-H2B-EGFP vector (Addgene Plasmid #25999) was used as a backbone. The H2B-EGFP coding sequence was removed with Hinc2 and Nsi1 and replaced with an EGFP-SspB-LARG coding sequence. SspB-LARG was PCR-generated from mCherry-NES-SspB-LARG-DH-P2A-iLIDcaax (Addgene Plasmid #173870) and EGFP was PCR-generated from the parental LV-H2B-EGFP plasmid. A six amino acid linker (GTSAGG) was introduced between EGFP and SspB. The PINCH-mEmerald (#54230, Addgene) vector was a kind gift from Michael Davidson. The Paxillin-EGFP (#15233, Addgene) vector was a kind gift from Rick Horwitz. The pLV-Stargazin-mTurquoise2-iLID vector (#161001, Addgene) was a kind gift from Sean Collins. The E-Cadherin-EGFP and myosin-RLC-iRFP vectors were a kind gift from Jordan Beach’s lab (Loyola University Chicago, Maywood, IL). The EGFP-VASP vector was a kind gift from Mary Beckerle (University of Utah, Salt Lake City, UT). The vinculin tension sensor vector (pRRL-vinTS) was a kind gift from Brenton Hoffman (Duke University, Durham, NC). The EGFP-Kindlin2 vector was a kind gift from Edward Plow (Cleveland Clinic, Cleveland, OH). The Zyxin-EGFP vector was a kind gift from Clare Waterman (NIH, Bethesda, MD). The Halo-Trip6 (VB240427-1344eus) and Halo-LIMD1 (VB240427-1342qgn) lentiviral vectors were purchased from Vectorbuilder. The vinculin-mEmerald and mCherry-vinculin vectors were a kind gift from Valerie Weaver (University of California San Francisco, San Francisco, CA). All constructs were verified by sequencing.

### Cell culture and generation of stable lines

Human foreskin fibroblasts (HFFs) were obtained from ATCC (CRL-2522). Mouse embryonic fibroblasts (MEFs), zyxin KO MEFs and zyxin KO MEFs stably re-expressing GFP-zyxin (zyxin-GFP MEFs) were a kind gift from Mary Beckerle (University of Utah, Salt Lake City, UT). Human breast epithelial cells (MCF10A) were a kind gift from Sean Fanning (Loyola University Chicago, Maywood, IL).

All stable lines were generated by lentiviral transduction using a transfer vector encoding the protein of interest, along with the psPAX2 packaging vector (#12260, Addgene) and the pMD2.G envelope vector (#12259, Addgene). MCF10A lines stably expressing Halo-TRIP6 and Halo-LIMD1 were generated using the VB240427-1344eus and VB240427-1342qgn lentiviral vectors, respectively. Zyxin KO MEFs and GFP-zyxin MEFs stably expressing myosin-RLC-iRFP670 were generated using the myosin-RLC-iRFP670 vector. HFFs stably expressing Stargazin-mTurquoise2-iLID were generated using the pLV-Stargazin-mTurquoise2-iLID vector.

To generate MEFs stably expressing vinTS, endogenous vinculin was first knocked down by lentiviral transduction of two shRNA constructs (TRCN0000295656 and TRCN0000288326, Sigma), followed by re-expression of vinTS via a second lentiviral transducton of the pRRL-vinTS vector.

The HFFs and MEF lines were cultured in DMEM (#10-013-CV, Corning) supplemented with 10% fetal bovine serum (#10437-028, Gibco) and 1% antibiotic-antimycotic solution (#30-004-Cl, Corning) at 37°C and 5% CO2. MCF10As were cultured in DMEM-F12 (#11330-032, Gibco) supplemented with 5% fetal bovine serum (#10437-028, Gibco), 20 ng/ml human epidermal growth factor (#AF-100-15-500UG, Peprotech), 0.5 *µ*g/ml Hydrocortisone (H-0888, Sigma), 100 ng/ml Cholera Toxin (C-8052, Sigma), 10 *µ*g/ml human Insulin (12585-014, Gibco) and 1% antibiotic-antimycotic solution (#30-004-Cl, Corning) at 37°C and 5% CO2.

### Transfection and labeling

At 24 h before each experiment, cells were transfected with 5 *µ*g total DNA using a Neon electroporation system (ThermoFisher Scientific) and plated on glass coverslips. For traction force microscopy experiments, cells were plated on polyacrylamide gels (see details below). For actin and Halo labeling, cells were incubated for 1 hour with 1 *µ*M Spy-Actin (CY-SC202, Cytoskeleton) or SiR-Actin (CY-SC001, Cytoskeleton), or overnight with 20 nM Halo dye (JFx650-HaloTag ligand, Lavis Lab, Lot: Sep-1-152), respectively.

### Live cell imaging

Cells were imaged in culture media at 37°C in 5% CO_2_. Live cell imaging was performed on a spinning disk confocal microscope from 3i (Intelligent Imaging Innovations) consisting of an Axio Observer 7 inverted microscope (Zeiss) attached to a W1 Confocal Spinning Disk (Yokogawa) with Mesa field flattening (Intelligent Imaging Innovations), a motorized X,Y stage (ASI), a Phasor photomanipulation unit (Intelligent Imaging Innovations) and a Prime 95B sCMOS (Photometrics) camera. Illumination was provided by a TTL triggered multifiber laser launch (Intelligent Imaging Innovations) consisting of six diode laser lines (405/445/488/514/561/640 nm) and all matching requisite filters, using a 63x 1.4 NA Plan-Apochromat objective (Zeiss). Temperature and humidity were maintained using a Bold Line full enclosure incubator (Oko Labs). The microscope was controlled using the Slidebook 6 Software (Intelligent Imaging Innovations). For FRET imaging, the vinTS construct was illuminated with the 445 nm laser and light was collected with a Semrock FF01-571/72 filter

### Adhesion intensity analysis

For photoablation experiments, a 5 *µ*m line was drawn in Slidebook 6 across the SF of interest. After steady state imaging, the marked region was illuminated with a 405 laser for 1.5 s at 400-600 *µ*W (glass) or 1.8-2.5 mW (gels) to reduce SF tension and by extension tension on adjacent FAs. To reduce tension at cell-cell junctions, two 5 *µ*m lines were drawn across adjacent SFs on each side of the junction illuminated with a 405 laser at 1.3-1.8 mW for 5 s. For FRAP experiments, marked regions were illuminated with a 488 laser at 150-250 *µ*W for 3 s. Imaging resumed immediately after photoablation/FRAP. Details on photoablation/FRAP timepoints, time lapse duration and cell counts are provided in the figures.

All movies were flatfield and photobleach corrected and registered prior to analysis as described previously [32]. For FRET analysis, movies were flatfield corrected and registered, but not photobleach corrected. Masks were created of the FA based on the fluorescence intensity and traction stress regions based on the traction maps. For cell-cell adhesions, a 7 × 9.5 *µ*m mask was drawn around the junction to account for movement of the adhesion.

Based on the masked regions, mean fluorescence intensity or traction stress was calculated for each timepoint, normalized to the mean pre-ablation state value. Traces from multiple movies were averaged and plotted as mean ± SD. Fluorescence intensity and traction stress changes were calculated by comparing the mean pre- and final ablation values for each movie, and plotted as boxplots for statistical analysis. Boxplots in Fig. 1F show the distribution of the peak mean FRET intensity after Y-27632 or media treatment, normalized to the mean pre-treatment values. Details on frame numbers (time intervals) used for normalization and statistical analysis are provided in the figure legends.

Kymographs in Fig. 5C-D and Fig. 2 were analyzed by calculating the sum intensity of each column (timepoint), normalized to the first 5 columns. Traces from multiple kymographs were then averaged and plotted as mean ± SD.

### Optogenetics

Before each time lapse, a ~ 27*µm*^2^ box was drawn across the SFs of interest in SlideBook 6. After imaging the steady state for 5 min, the marked region was illuminated with a 405 nm laser for 10 min at 4.5 *µ*W to locally increase SF tension, followed by an additional 5 min of imaging. Cell counts are provided in the figures.

Images were registered, flat-field and photobleach corrected. FA masks were generated to quantify and plot fluorescence intensity versus traction stresses within regions where traction stresses increased, remained stable, or decreased. In Fig. 1H and Fig. 2G, each point represents the mean fluorescence intensity and traction stress over the last 100 s of each time lapse, normalized to the 100 s before activation. In Fig. 1A-B, each point represents the mean fluorescence intensity and traction stress at each time point, normalized to the 100 s before activation.

### Actin retrograde flow analysis

zyxin-GFP MEFs and zyxin KO MEFs (expressing mEmerald-vinculin) were imaged every 5 s for 5 min. Both lines stably expressed myosin-RLC-iRFP670, which was used for retrograde flow analysis due to its striated pattern along SFs. Movies were registered based on the zyxin-GFP or mEmerald-vinculin channels. Only FA-coupled SFs were analyzed. Kymographs were drawn across the FA-SF interface based on the zyxin/vinculin (FA) and actin (SF) images. Retrograde flow velocity (nm/s) was calculated based on the myosin striations in the first 100 s of each movie. Multiple regions were analyzed per cell when possible (zyxin-GFP MEFs: 15 cells, 37 regions; zyxin KO MEFs: 17 cells, 38 regions), and the mean flow velocity per cell was plotted as boxplots and used for statistical analysis.

### Traction force microscopy

Traction force microscopy was performed as described previously [32, 34]. Coverslips were prepared by incubating with a 2% solution of 3-aminopropyltrimethyoxysilane (313255000, Acros Organics) diluted in isopropanol, followed by fixation in 1% glutaraldehyde (16360, Electron Microscopy Sciences) in ddH20. Polyacrylamide gels (shear modulus: 16 kPa, final concentrations of 12% acrylamide [1610140, Bio-Rad] and 0.15% bisacrylamide [1610142, Bio-Rad]) were mixed with 0.04 *µ*m fluorescent microspheres (F8789, Invitrogen) and polymerized on activated glass coverslips for 1 h at room temperature. After polymerization, gels were rehydrated for 45 min and coupled to 50 *µ*L of 1 mg/ml human plasma fibronectin (FC010, Millipore) for 1 h at room temperature using the photoactivatable cross-linker Sulfo-Sanpah (22589, Pierce Scientific). Following fibronectin cross-linking, cells were plated on the gels and allowed to spread overnight. The next day, images were taken of the cells and underlying fluorescent beads.

Following imaging, cells were removed from the gel using 0.05% SDS and a reference image of the fluorescent beads in the unstrained gel was taken. Analysis of traction forces was performed using code written in Python according to previously described approaches [34]. Prior to processing, images were flat-field corrected and the reference bead image was aligned to the bead image with the cell attached. Displacements in the beads were calculated using an optical flow algorithm in OpenCV (Open Source Computer Vision Library, https://github/itseez/opencv) with a window size of 8 pixels. Traction stresses were calculated using the Fourier Transform Traction Cytometry approach with a regularization parameter of 1 × 10^*−*7^.

### Statistical analysis

Statistical analyses were performed in Python using the non-parametric multiple comparison Kruskal-Wallis test with Dunn’s post-hoc test. P values < 0.05 were considered statistically significant.

### Software

All images were exported from Slidebook as 16-bit TIFF files and analyzed in Python and Fiji. All custom code, including traction force microscopy routines, are available at (https://github.com/OakesLab).

## Acknowledgements

We thank the other members of the Beach and Oakes labs for many helpful discussions. We are also thankful to the Waterman, Weaver, Beckerle, Hoffman and Plow and Horwitz labs for providing us with vectors and cell lines used in this study. This work was supported by NIH NIGMS grant R01-GM148644 to P.W.O.

## Author contributions

SS and PWO conceived the study. SS and SC performed experiments. LT, HW, and JRB contributed reagents and helped develop assays. SS, SC, and PWO performed data analysis. SS and PWO wrote the manuscript.

