## Supplemental Figures for "LIM Domain Proteins link molecular and global tension by recognizing strained actin in adhesions"

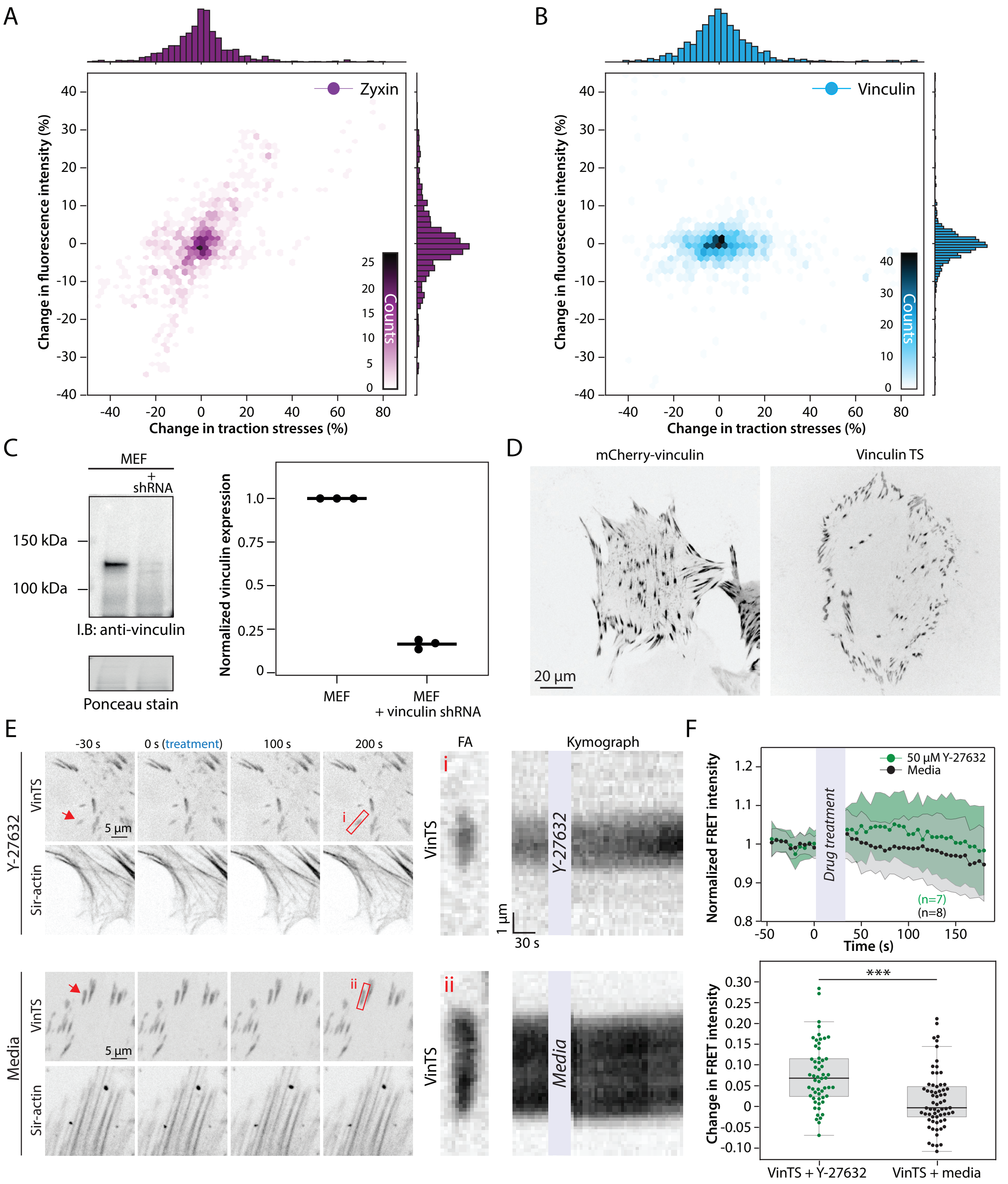


**Fig. S 1. Vinculin FA localization is unresponsive to local changes in adhesion tension while its intramolecular tension decreases upon global cytoskeletal remodeling. A-B)** Distribution of normalized mApple-zyxin (A) and mCherry-vinculin (B) intensities vs. traction stresses in focal adhesions across all timepoints from optogenetic experiments in Fig. 1F-H (zyxin) and Fig. 2E-G (vinculin). **C)** Western blot analysis of endogenous vinculin levels in MEFs with and without stable vinculin shRNA expression. **D)** MEFs expressing vinculin-mCherry (left) or vinculin knockdown MEFs exressing vinTS (right), showing focal adhesion localization of both constructs. **E)** Vinculin knockdown MEFs expressing vinTS, treated with 50 $\mu$M Y27632 or media. Red arrows mark the adhesions used for analysis Red boxes indicate FAs used to generate kymographs. **F)** Normalized FRET intensity $\pm$ SD. Boxplots show change in average FRET intensity between the pre-treatment and peak intensity of each FA analyzed. n values indicate the number of cells analyzed.

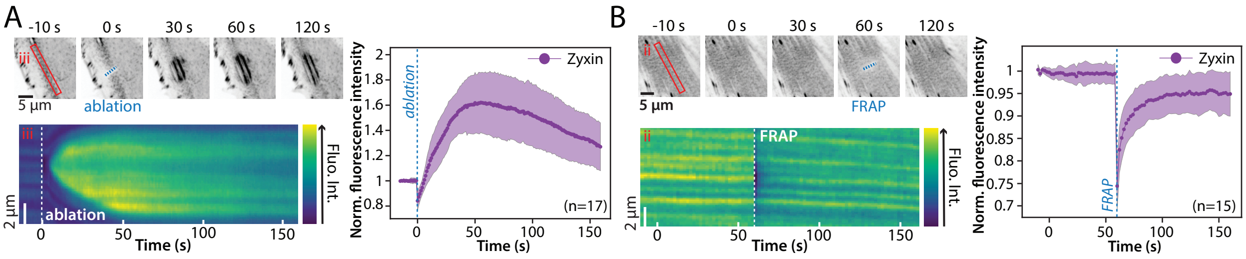


**Fig. S2. Zyxin response to photoablation and FRAP in SFs. A)** Representative images and kymographs (red boxes) of MEFs expressing eGFP-zyxin during a photoablation (blue dashed line). Plot shows the normalized zyxin intensity over time in the strain site. **B)** Representative images and kymographs (red boxes) of MEFs expressing eGFP-zyxin during FRAP (blue dashed line) of a SF. Plot shows the normalized zyxin intensity over time in the SF.

### Supplemental Movies

**Supplemental Movie 1.** Movie 1: Photoablation of SFs reduces tension on attached FAs, leading to decreased zyxin localization and traction stresses at the adhesions (Fig. 1A)

**Supplemental Movie 2.** Movie 2: Optogenetic modulation of adhesion tension reveals a positive correlation between local traction stress magnitude and zyxin intensity (Fig. 2G), but not vinculin intensity (Fig. 2F), at adhesions

**Supplemental Movie 3.** Movie 3: Photoablation of SFs lowers tension on attached FAs, causing a decrease in zyxin localization, but not in vinculin localization, at these sites (Fig. 2B)

**Supplemental Movie 4.** Movie 4: Vinculin intramolecular tension remains unchanged upon reducing adhesion tension via SF photoablation (Fig. 2I)

**Supplemental Movie 5.** Movie 5: FA localization of paxillin decreases upon reducing adhesion tension via SF photoablation, while PINCH and kindlin2 intensities remain unchanged (Fig. 3A-C)

**Supplemental Movie 6.** Movie 6: Cell-cell junction localization of Trip6 and LIMD1, but not E-cadherin, decreases upon reducing adhesion tension via SF photoablation (Fig. 3D-F)

**Supplemental Movie 7.** Movie 7: VASP localization at FAs is reduced in WT, but not zyxin KO, MEFs upon reducing adhesion tension via SF photoablation (Fig. 4A)
